# Low numbers of cytokine transcripts drive inflammatory skin diseases by initiating amplification cascades in localized epidermal clusters

**DOI:** 10.1101/2021.06.10.447894

**Authors:** A. Schäbitz, C. Hillig, A. Farnoud, M. Jargosch, E. Scala, A.C. Pilz, N. Bhalla, M. Mubarak, J. Thomas, M. Stahle, T. Biedermann, C.B. Schmidt-Weber, F. Theis, N. Garzorz-Stark, K. Eyerich K, M.P. Menden, S. Eyerich

**Affiliations:** Division of Dermatology and Venereology, Department of Medicine Solna, and Center for Molecular Medicine, Karolinska Institutet, Stockholm, Sweden; Institute of Computational Biology, Helmholtz Zentrum München - German Research Centre for Environmental Health, Ingolstädter Landstrasse 1, 85764 Neuherberg, Germany; Center for Allergy and Environment (ZAUM), Technical University and Helmholtz Center Munich, Biedersteinerstrasse 29, 80802 Munich, Germany; Department of Dermatology and Allergy, Technical University of Munich, Biedersteinerstrasse 29, 80802 Munich, Germany; Department of Gene Technology, School of Engineering Sciences in Chemistry, Biotechnology and Health, KTH Royal Institute of Technology, Stockholm, Sweden; Department of Dermatology and Venereology, Unit of Dermatology, Karolinska University Hospital, Stockholm, Sweden; Department of Biology, Ludwig-Maximilians University, Goßhadernerstrasse 2, Martinsried, 82152, Germany; German Center for Diabetes Research (DZD e.V.), Ingolstädter Landstrasse 1, 85764 Neuherberg, Germany

**Keywords:** spatial transcriptomics, chronic inflammatory skin disease, cytokines, skin, psoriasis, lichen, atopic dermatitis

## Abstract

Abundant polyclonal T cells infiltrate chronic inflammatory diseases and characterization of these cells is needed to distinguish disease-driving from bystander immune cells. Here, we investigated 52,000 human cutaneous transcriptomes of non-communicable inflammatory skin diseases (ncISD) using spatial transcriptomics. Despite the expected T cell infiltration, we observed only 1-10 pathogenic T cell cytokine per skin section. Cytokine expression was limited to lesional skin and presented in a disease-specific pattern. In fact, we identified responder signatures in direct proximity of cytokines, and showed that single cytokine transcripts initiate amplification cascades of thousands of specific responder transcripts forming localized epidermal clusters. Thus, within the abundant and polyclonal T cell infiltrates of ncISD, only a few T cells drive disease by initiating an inflammatory amplification cascade in their local microenvironment.

## Introduction

Non-communicable inflammatory diseases are based on complex interactions of predisposing genetic background and environmental triggers that collectively result in altered immune responses. Several hundred non-communicable inflammatory skin diseases (ncISD) exist, including psoriasis, atopic dermatitis (AD), and lichen planus (lichen). Despite their heterogeneity, most ncISD can be categorised according to adaptive immune pathways based on the interaction of distinct lymphocyte subsets with the epithelium(*1, 2*). Whereas psoriasis represents a classical type 3 immune cell mediated disease(*3, 4*), AD is dominated by type 2(*5, 6*), and lichen by type 1(*7, 8*) immune cells. Accordingly, psoriasis can be efficiently treated with antibodies targeting cytokines of type 3 immunity, i.e., IL-17A or IL-23(*9, 10*). Likewise, AD is successfully treated with antibodies targeting cytokines of type 2 immune cells, such as IL-13(*11, 12*). However, without models to predict therapeutic responses, many patients do not respond to a given therapy. Furthermore, we lack curative approaches, since current therapies neutralize cytokines, but do not target antigen-specificity. More granular information regarding the profile, kinetics, and spatial distribution of cytokine-secreting immune cells is needed to achieve a substantial advance in addressing these challenges.

Emerging molecular techniques allow analysis of mRNA expression in single-cell and spatial contexts, thus enabling deep phenotyping of relevant cell types in ncISD(*13, 14*). Conventional single-cell sequencing techniques require dissociation of the tissue and thereby might bias the interpretation due to loss of tissue context. Spatial transcriptomics (ST) overcomes this issue, allowing the study of the inflamed skin architecture(*15, 16*), but does not provide single cell resolution. Investigating disease-driving cells together with their direct responder signatures in a spatial context will offer new insights into the pathogenic microenvironment of ncISD.

Here, we investigated adaptive immune responses in lesional and non-lesional skin of ncISD with spatial resolution. We observed that single transcripts of disease-driving T cell cytokines, namely *IL17A* for psoriasis, *IL13* for AD, and *IFNG* for lichen planus, initiated localized amplification cascades of specific inflammatory responder genes that collectively represent hallmarks of the respective disease pathogenesis. Thus, a few T cells drive ncISD within an abundant polyclonal infiltrate.

## Results

To analyze the pathogenic microenvironment across multiple ST sections of non-lesional and lesional ncISD skin, we characterised the spatial transcriptomic landscape of ncISD (Fig. 1A), covering psoriasis, AD, lichen, and pityriasis rubra pilaris (PRP). This dataset included 64 samples (18 lesional, 14 non-lesional in duplicates) and the transcriptomes of 52,020 spots. After removing 8,377 spots with low unique molecular identifier (UMI) counts and high mitochondrial fraction, 15,285 non-lesional and 28,358 lesional spots entered further analyses.

**Figure 1:**
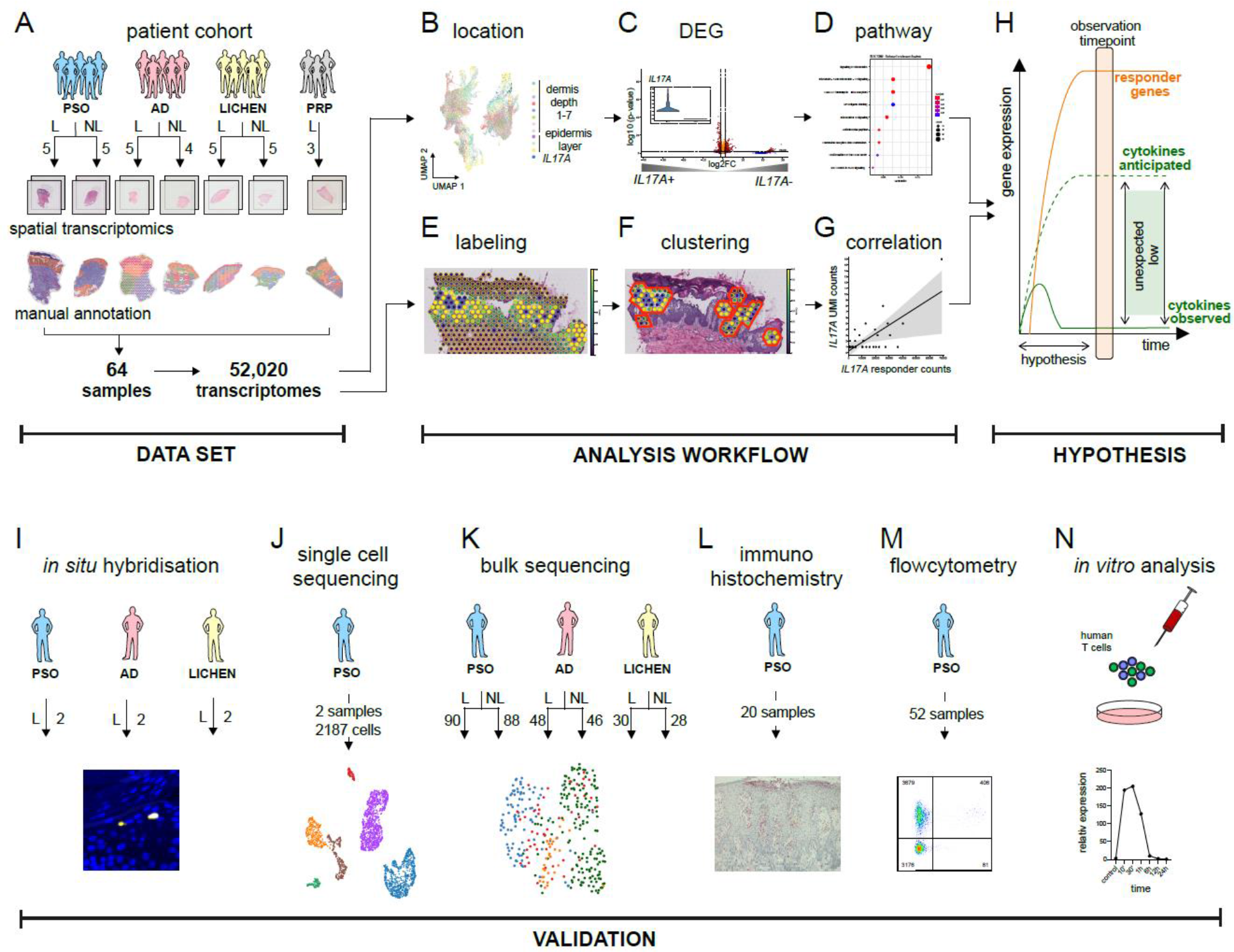
The study design highlighting the spatial transcriptomic data set, the analysis pipeline, and the validation cohorts and techniques. **A)** ST dataset consisting of 64 spatial samples (18 patients, 36 lesional samples, 28 non-lesional samples, and four ncISD (psoriasis, atopic dermatitis (AD), lichen planus and pityriasis rubra pilaris (PRP)) resulting in 52,020 transcriptomes. Every spot in all samples was manually annotated according to tissue localization (basal-, middle-, upper epidermis, dermis 1-7). Analysis workflow including **B)** the assignment of each spatial spot to a tissue localization, **C)** differential gene expression (DEG) analysis of cytokine-positive versus cytokine-negative spots, and **D)** pathway analysis. For spatial correlation of cytokine-positive spots with cytokine responder genes: **E)** spots were labeled as cytokine or responder positive, **F)** clusters of cytokines and responders were defined, and **G)** correlation analysis was performed. **H)** Hypothesis expecting higher cytokine mRNA counts than observed. Low cytokine counts in ncISD were confirmed using **I)** *in situ* hybridization, **J)** single cell sequencing, **K)** bulk sequencing, **L)** immunohistochemistry, **M)** flow cytometry, and **N)** *in vitro* stimulation of human T cells

We proposed two complementary analysis workflows. The first workflow incorporated spatial features in differential gene expression (DEG) analysis of spots containing cytokine-positive *versus* cytokine-negative leukocytes, followed by pathway enrichment analyses (Fig. 1B-D; Methods). The second workflow labelled cytokine-positive spots, and then used a density-based clustering method to boost correlations of cytokine and responder gene signatures according to spatial features (Fig. 1E-G; Methods). This analysis led to the surprising observation that single cytokine transcripts initiated amplification cascades of thousands of specific responder transcripts, which are causative and disease driving in the tissue micro-environment (Fig. 1H). We validated the results using a variety of patient cohorts and techniques such as *in situ* hybridisation, single-cell and bulk sequencing, immunohistochemistry, flow cytometry and cell culture analysis (Fig. 1I-N).

### Low numbers of disease-driving cytokine transcripts are expressed in lesional skin of ncISD

The tissue inflammation of ncISD is driven by T cell cytokines, therefore we examined the expression of the major effector cytokines driving the common ncISD psoriasis, AD, and lichen, namely *IL17A*, *IFNG*, and *IL13*, respectively (Fig. 2A). As expected, the number of cytokine UMI counts was low in non-lesional skin samples. However, even in lesional ncISD skin, we detected only a few cytokine transcripts (Fig. 2B-E). The spatial distribution, however, was distinct for the investigated cytokines. *IL17A* was detected in all layers of the lesional epidermis and was virtually absent in the dermis (epidermis vs dermis p=2.96e^−13^), while *IFNG* (epidermis + dermis 1 vs dermis 2-7 p=1.17e^−37^) and *IL13* (basal epidermis + dermis 1 vs upper + middle epidermis + dermis 2-7 p=0.0016) were mostly expressed in the basal epidermis and upper dermis layers (Fig. 2A, C, Fig. S1A). Taking the whole section into account, we detected only a few transcripts for *IFNG, IL13* or *IL17A* (272, 57, or 92 UMI counts in all sections respectively) in lesional skin (Fig. 2E).

**Figure 2:**
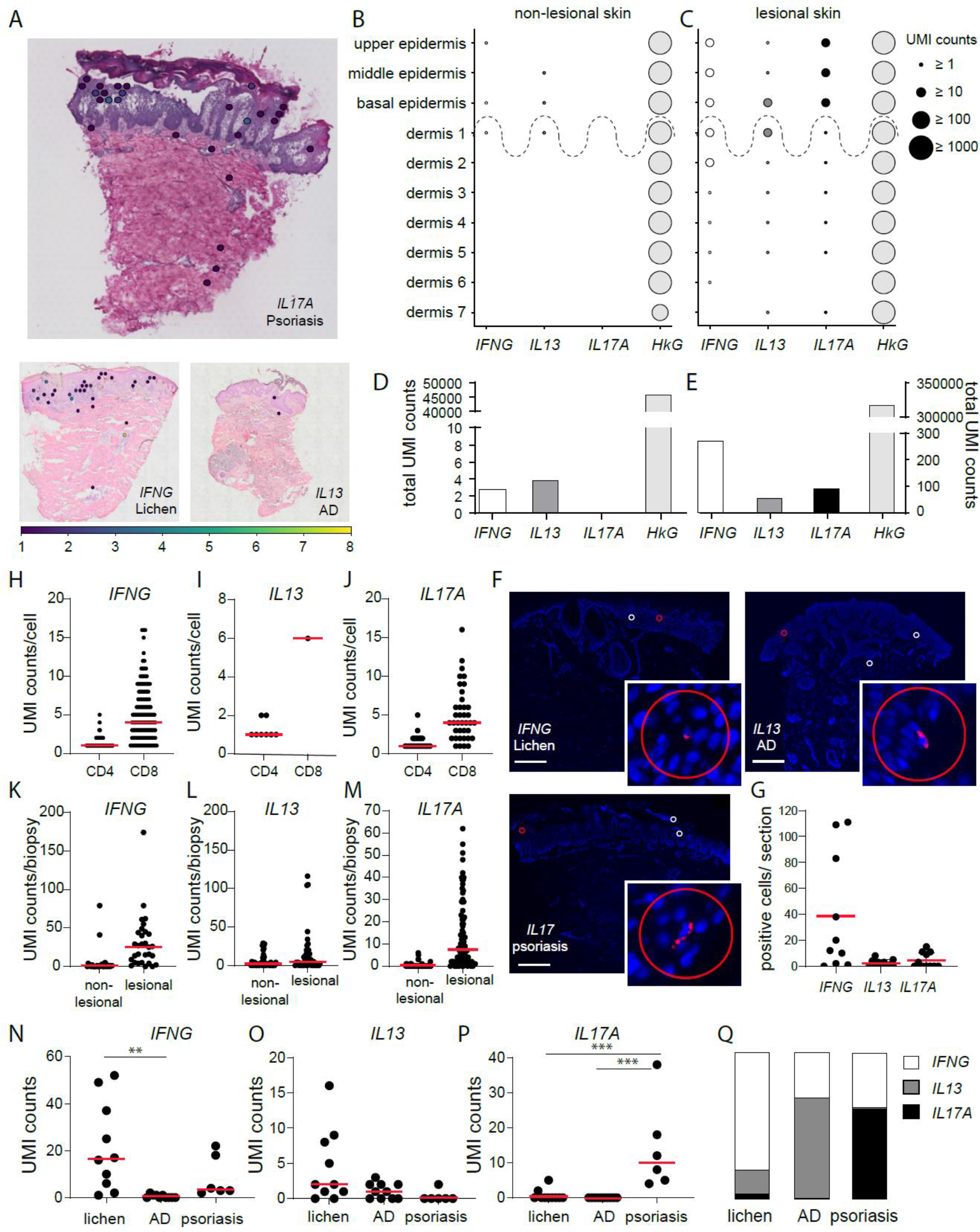
Low numbers of disease-driving cytokine transcripts are expressed in lesional skin of ncISD. **A)** Representative ST sections for psoriasis with *IL17A*+ spots, AD with *IL13*+ spots, and lichen with *IFNG*+ spots. **B, C)** UMI-counts of *IFNG, IL13* and *IL17A* expressed in the manually annotated tissue layers ‘upper, middle, and basal epidermis’ and ‘dermis depth 1-7’ in non-lesional and lesional skin of all investigated samples (n=56). *GAPDH* serves as a housekeeping gene (HkG). **D, E)** Total cytokine and *GAPDH* UMI counts in all non-lesional (D) and lesional (E) skin sections. **F)** *In situ* hybridization for *IFNG, IL13* and *IL17A* in representative stainings of lichen (upper left panel), AD (upper right panel) and psoriasis (lower panel). Scale bar indicates 500 μm; circles represent Ø 55 μm. **G)** Quantification of cytokine positive cells per *in situ* section. **H-J)** scRNA-seq analysis of psoriasis biopsies (n=2, 2,187 cells) indicating the UMI count of *IFNG* (178 cells), *IL13* (9 cells), and *IL17A* (61 cells) per cell in CD4 or CD8 co-expressing cells. **K-M)** Bulk sequencing analysis of non-lesional and lesional lichen (n=30) *(IFNG*), AD (n=48) (*IL13*), and psoriasis (n=90) (*IL17A*) biopsies indicating the total UMI counts for *IFNG, IL13*, and *IL17A*, respectively, in each biopsy. **N-P)** UMI counts for *IFNG, IL13*, and *IL17A* in ST sections separated by disease (each dot represents one section). **Q)** Percentage of disease relevant cytokines in lichen, AD, and psoriasis normalised to 100%.

We validated the low transcript numbers and low numbers of cytokine-positive cells in inflamed tissue using various *ex vivo* and *in vitro* methods. *In situ* hybridization identified very few cytokine-positive signals (Fig. 2F). The median number of positive cells per section for *IL17A*, *IFNG*, and *IL13* mRNA were 0, 16, and 2 for psoriasis, AD, and lichen, respectively, thus confirming our observations from the ST analysis (Fig. 2G). Single-cell RNASeq analysis of psoriasis also indicated few transcripts per *IL17A+* or *IFNG+* cell, respectively, with a median UMI count for *IL17A* or *IFNG* of 1/ CD4+ cell and 4/ CD8+ cell (Fig. 2H-J). We also investigated a large cohort of ncISD patients using bulk RNA sequencing. Here, in a third of a 6 mm skin punch biopsy we detected a median of 0 and 7.5 *IL17A* transcripts/biopsy in non-lesional and lesional psoriasis skin, respectively (Fig. 2K-M). AD presented with a median *IL13* UMI count of 2 and 4,5/biopsy and lichen with a median *IFNG* UMI count of 1 and 25,5/biopsy in non-lesional and lesional skin, respectively. Immunohistochemistry and flow cytometric analysis of skin-infiltrating T cells revealed comparable numbers of cytokine-positive lymphocytes in lesional skin (histology: 13.3% IL-17A+ lymphocytes, flow cytometry: 4.2% CD4+IL-17A+, 4.9% CD8+IL-17A+ (Fig. S5A, B, C). Time course analysis showed that short T cell receptor (TCR) stimulation *in vitro* resulted in transient mRNA production with a peak at 10-30 minutes and a total production time of less than 6 hours. Low numbers of mRNA transcripts per cell increased with prolonged TCR stimulation (Fig. S5D).

Despite their low UMI counts, cytokines showed a disease-specific expression pattern. *IL17A* transcripts were mostly expressed in lesional psoriasis, *IFNG* in lichen, and *IL13* in lichen and AD (Fig. 2N-P, Fig. S2A, C) with an emphasized expression in upper skin layers (Fig. 3A, Fig. S2B, C). This held true for other disease-driving T cell cytokines such as *IL17F, IL21, IL22, TNFA, IL10*, and *IL4* (Fig. S1B). The relative distribution of the signature cytokines confirmed that psoriasis is a type 3, AD a type 2, and lichen a type 1 immune-driven disease (Fig. 2Q, Figure S1B, C). Taken together, these findings show that low numbers of disease-specific cytokine transcripts are produced by a few T cells that have a characteristic tissue distribution.

**Figure 3:**
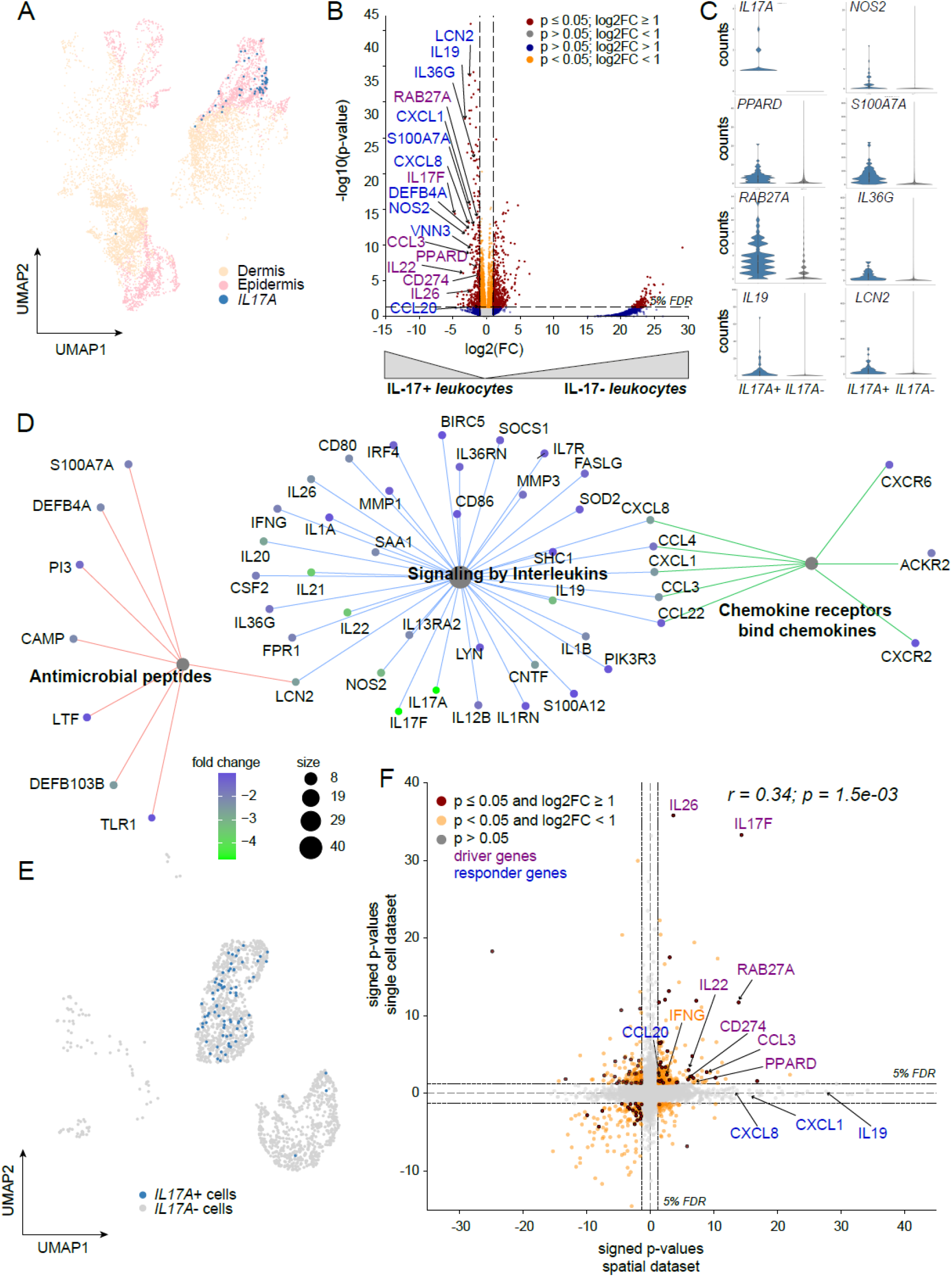
IL17A transcript positive spots are characterized by Th17 markers and IL-17A tissue response genes. **A)** ST data spots expressing leukocyte marker genes and their location in epidermis or dermis. *IL17A* positive spots are highlighted in blue. **B)** Volcano plot analysing the gene expression profile of *IL17A* positive (*IL17A*+) *versus IL17* negative (*IL17A*−) spots. Purple labeling indicates genes expressed in leukocytes; blue labeling indicates response genes of *IL17A* in the skin. Coordinates for *IL17A* (−37.7/89.2) are not shown. **C)** Violin plots of selected genes in *IL17A*+ and *IL17A*− spots indicate the general low expression of gold standard genes. **D)** Pathway enrichment analysis of genes co-expressed with *IL17A* in spatial spots. Here, the log2FC score ranges between 0.0 and −4.7 highlighting *IL-17A* (log2FC= −37.7) in the same colour as *IL-17F* (log2FC=−4.7). **E)** scRNA-seq analysis of psoriasis skin highlighting *IL17A* expression in lymphocytes (blue) in a UMAP (Uniform Manifold Approximation and Projection) plot. **F)** Comparison of the DEG analysis of *IL17A* positive spatial spots and *IL17A* positive single cells in a signed p-value plot. Significantly expressed genes in both data sets are shown in red. Purple and blue labeling indicates T-cell derived genes and skin response genes, respectively. *IFNG* is marked in orange.

### Cytokine-producing T cells and nearby cells are characterized by specific driver and responder gene signatures

To specifically phenotype these cytokine-producing cells, we performed DEG analysis of spots containing cytokine-positive leukocytes compared to leukocyte spots without cytokine expression. DEGs generally consisted of T cell genes and genes induced by the cytokines in cells in close proximity, so-called responder genes. T cell genes associated with *IL17A* were *IL17F, IL22*, and *IL26*, and the responder signature of *IL17A* consisted of e.g., *IL19, NOS2, S100A7A, DEFB4A, CXCL8*, and *IL36G* (Fig. 3B, C). *IFNG* positive spots were characterized by genes related to type 1 immune cells, such as *GZMB, FASLG, CD70, CXCR3*, and *CXCR6*, and by *IFNG*-dependent response genes such as *CXCL9, CXCL10, CXCL11*, and *CXCL13* (Fig. S3A). *IL13* positive spots presented themselves with differentially expressed genes associated with type 2 cells, such as *IL2, IL10*, and *CD48*, plus genes associated with their response, among them *CCL17, CCL19, CCL26*, and *OSM* (Fig. S3C). This strength of ST in revealing driver genes together with their correlating responders was further illustrated by apathway enrichment analysis of lead cytokine-positive spots, showing specific signatures for both inflammation-driven cell signaling and tissue reaction to inflammation (Figure 3D, Figure S3B, D).

The identified immune cell driver genes were confirmed by our psoriasis scRNA-seq dataset. Here, unsupervised clustering identified distinct cell types (Fig. S4A), and disease-driving cytokines were exclusively detected in the leukocyte and antigen-presenting cell cluster (Fig. 3E, Fig. S4B). Confirming the spatial dataset, most leukocyte-associated genes were also identified in the single-cell data, whereas the responder gene signatures were widely missing in the single-cell data (Fig. 3F, Fig. S4C, D).

Besides identifying gene signatures specific for cytokine-producing T cells in their local microenvironment, we established a gene signature of 14 genes by combining the DEG lists of *IL17A, IFNG*, and *IL13* to identify genes associated with general T cell activation. This signature was comprised of cytokines (*IFNG, IL-22, CSF2*, *IL19*), chemokines (*CXCR6, CCL3*), and further markers of T cell activation (*CD80, GZMB, LILRB3, TNFRSF9, FUT7, NR4A3, G0S2, LAG3*) (Fig. S3F).

In essence, we identified gene signatures that define cytokine-producing T cells in lesional skin as well as responder signatures of genes induced in close spatial proximity by these cytokines in the inflammatory microenvironment.

### Immune response is spatially correlated with cytokine transcript number

To investigate the functional relevance of the few cytokine transcripts in lesional ncISD skin, we studied the correlation between cytokine-secreting cells and their responder signatures (Fig. 1). To further corroborate the epithelial response signatures for *IL17A, IFNG*, and *IL13*, we compared the specific expression pattern of primary human keratinocytes that were stimulated with recombinant IL-17A, IFN-γ, or IL-13 *in vitro* with the spatial DEGs for each cytokine. Genes that were present in both datasets were used as cytokine-specific tissue responder genes (Fig. S6). Initially, we correlated these responder gene counts with their matching cytokine counts in all ST sections without taking the spatial resolution into account (Fig. 4A-C). *IL17A* and *IL13* had low positive correlations with the respective responder genes (Pearson r=0.37; p=4.8e^−3^ and r=0.42; p=1.21e^−3^, respectively), and *IFNG* had a strong correlation with its responders (Pearson r=0.78; p=9.94e^−13^). Correlations between cytokine counts and counts for responder genes of a different cytokine were either non-significant or lower (Fig. S7A-C). Conditional clustering of spatial information markedly improved the correlation between cytokines and tissue response in the inflammatory microenvironment (Fig. 4D-I, Methods). The responder maximum UMI counts showed a trend of being higher in close vicinity to cytokine-positive spots (Fig. 4D-F). The inclusion of the spatial information resulted in strong positive correlations for *IL17A* (Pearson r=0.91; p=9.61e^−13^), *IFNG* (Pearson r=0.85; p=1.56e^−23^), and *IL13* (Pearson r=0.71; p=9.56e^−4^) (Fig. 4G-I). We observed a strong weighted correlation between *IL17A* and signature responses of *IFNG* and *IL13* (Pearson r=0.95 and r=0.86, respectively) upon including the spatial information (Fig. S7D). This did not apply for the correlation of *IFNG* and *IL13* and the interchanged response. Here, the relationship strength was maintained (Fig. S7E-F).

**Figure 4:**
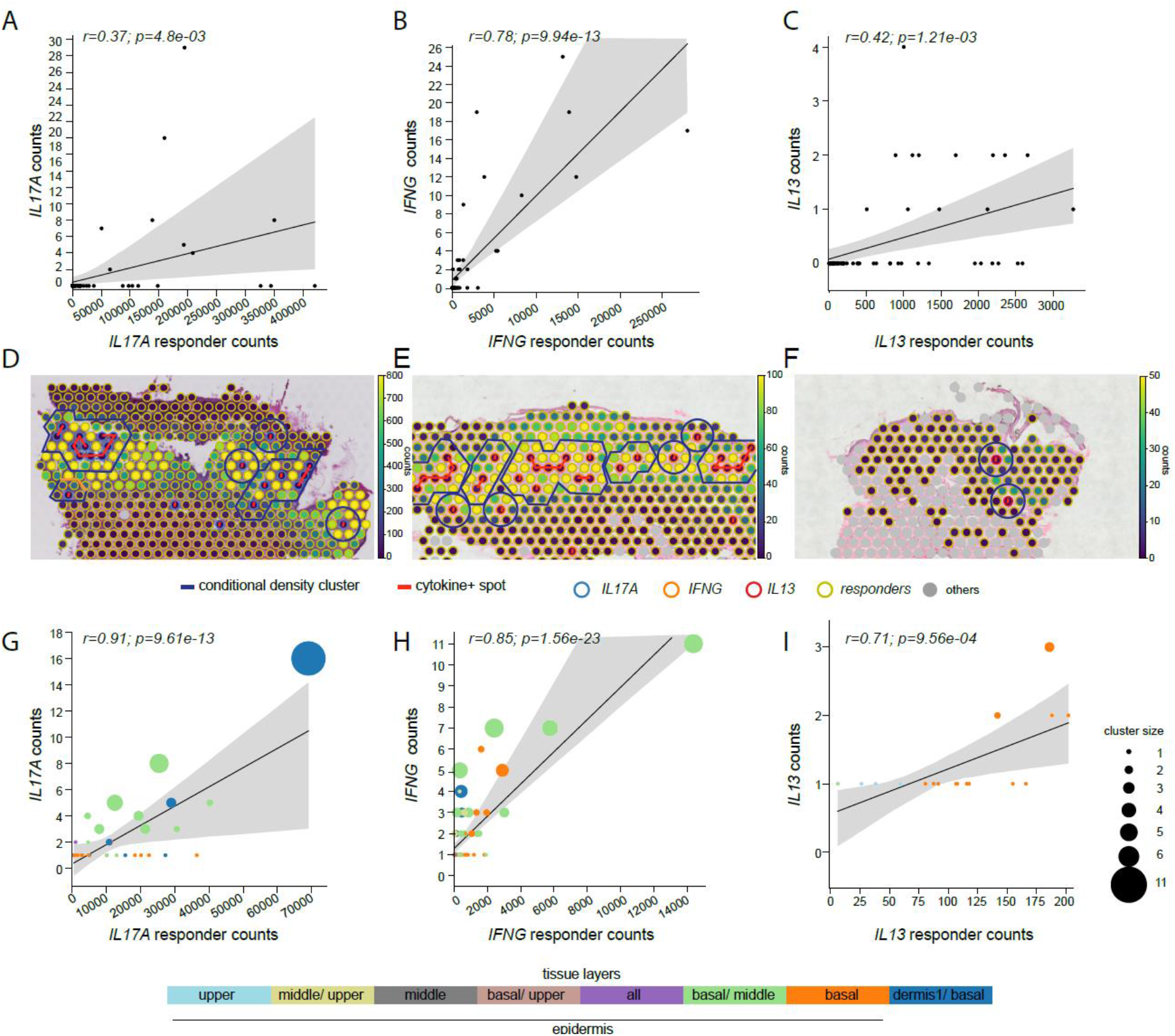
Immune response is spatially correlated with cytokine transcript number. **A-C)** Pearson correlation between cytokine positive and responder gene positive spots per tissue slice in the epidermis where each point in the plot represents the sum of all cytokine and responder counts in a tissue sample. **D-F)** Tissue slices showing the representative cytokines in relation to the responder signatures for psoriasis (D), lichen planus (E), and atopic dermatitis (F). Moreover, the counts of cytokines and responders on each spot are shown and the density-based clusters are highlighted in blue and graphs connecting cytokine positive spots are shown in red. **G-I)** Weighted spatial correlation incorporating the spatial relation of cytokines and their response located in the epidermis. Each point in the plots represents the sum of the counts of cytokines and responders in a cluster and the size of each point represents the number of cytokines in a cluster. The color of each spot is associated with the corresponding tissue layer.

In summary, regions with more cytokine transcripts had higher response signatures. Consequently, the inclusion of the spatial information and density-based clustering enhanced the biological signal for all cytokines and their response signatures. Altogether, these results provide comprehensive insights into the relationship between cytokine-expressing cells and their induced tissue response and confirm our hypothesis that a low number of transcripts is sufficient to induce pathogenic immune responses in the skin.

## Discussion

Causative therapies of common inflammatory skin diseases have seemed unrealistic as the diseases typically show an infiltrate of abundant and polyclonal immune cells into lesional skin. However, new molecular techniques and bioinformatic tools allow us to dissect ncISD on a new level and to undertake first steps in the development of causative therapies. Here, we investigated ncISD with spatial resolution. Namely, we explored the molecular landscape using DEG analysis and developed an algorithm to investigate the impact of cytokine-secreting cells on their direct surrounding environment. We demonstrated that a minority of immune cells actively drive the pathology of ncISD by producing low numbers of signature cytokine transcripts. Indeed, these few cytokine transcripts then translate into thousand-fold higher induction of pro-inflammatory response genes, thus forming an inflammatory microenvironment and subsequently leading to tissue damage.

Despite cytokine transcripts being rare in inflamed skin, they were detectable in disease- and spatial-specific patterns. The distribution matched that of antigens previously described in ncISD. In psoriasis, cytokine-secreting leukocytes were almost exclusively found throughout the epidermis, where epidermal and melanocytic autoantigens of psoriasis are expressed, e.g., ADAMTSL5(*17*), LL37(*18*), or lipid antigens presented via CD1(*12*). By contrast, antigens reported in lichen are located at the interface of the basal epidermis and the upper dermis, e.g., DSG(*19*), and several Hom s proteins(*20*) as potential antigens of AD are expressed in a similar location.

We characterized the cytokine-secreting cells in a tissue-dependent manner by implementing the annotations as a covariate. In the spatial context, we identified a reliable response signature of type 3 immune responses mediated by IL-17A, IL-17F, and IL-21 to be markers of oxidative stress such as NOS2, neutrophil migration such as CXCL8, and antimicrobial peptides like S100A7A and DEFB4A. By contrast, markers of type 1 immunity were chemokines such as CXCL9(*21*), CXCL10, and cytotoxic markers. The role for IFN-γ mediated apoptosis and necroptosis in type 1 ncISD is well established(*7, 8*) and is reflected by the expression of *FASL* and *GZMB* in *IFNG*+ spots. Type 2 immunity showed the least well-defined response signature, mostly built of type 2 attracting chemokines such as *CCL17, CCL19*, and *CCL26*. This signature was exclusively mediated by *IL13* as *IL4* transcripts were virtually undetectable in lesional skin even of AD.

The amplification cascade described here explains why response genes rather than the signature cytokines themselves are currently suggested as robust biomarkers for diagnostic or theranostic purposes in ncISD. Examples are a molecular classifier for differential diagnosis of psoriasis and eczema using *NOS2* and *CCL27*(*22, 23*), prediction of the response to anti-IL-17 therapies in psoriasis by IL-19 levels in serum(*24*), as well as correlation of the severity of psoriasis with DEFB4A(*25*) or the severity of AD with CCL17/TARC(*26*).

A reliable identification of disease-driving T cells and their cognate antigen might pave the way for causative treatment strategies of ncISD, e.g., antigen-specific immunotherapy. This has been attempted e.g., in AD as a global strategy with modest clinical efficacy(*27*), most likely because there is the need to identify disease endotypes defined by antigen-specificity of disease-driving T cells, within this heterogeneous disease. The proof-of-principle that causative therapies of ncISD are possible was made in the autoimmune blistering disease pemphigus vulgaris. Here, the causative antigen desmoglein 3 (DSG3) is identical in most patients and it is thus possible to design targeted therapies for the whole patient group. In fact, modified CAR T cell approaches neutralizing exclusively Dsg3-specific cells resulted in impressive and sustainable clinical improvements(*28, 29*).

Our conditional density-clustering method that extracts a correlation using the spatial information of the defined clusters based on cytokine-positive spots and their shared tissue-specific responders on in each tissue slice can be generalised to other diseases and tissues and represents a resource for identifying biomarkers and disease drivers. By integrating three-dimensional spatial information using consecutive tissue sections, the algorithm could be improved to identify disease-driving networks across tissue sections from the same patient, identifying antigen-specific T cell activation and promising a more precise treatment strategy.

A blueprint for successful precision medicine can be found in recent developments in oncology. Typically, tumors such as malignant melanoma are characterized by thousands of distinct mutations(*30*). However, few of them are actually driver mutations leading to tumor growth and metastasis(*31*). Targeting these driver mutations by specific targeted small molecules has led to dramatically increased survival rates of melanoma patients in recent years(*32*). Here, we demonstrate parallels to inflammatory skin diseases – non-cytokine-secreting T cells may be seen as irrelevant bystander cells, while targeting cytokine-secreting T cells is a promising strategy for effective and potentially causative treatments of ncISD. A prerequisite is to localize these cells in the inflammatory microenvironment and to identify the specific antigen that disease-driving T cells react against, which may pave the way for precision medicine in ncISD.

## Acknowledgements

The authors acknowledge support from the National Genomics Infrastructure in Stockholm funded by Science for Life Laboratory, the Knut and Alice Wallenberg Foundation and the Swedish Research Council, and SNIC/Uppsala Multidisciplinary Center for Advanced Computational Science for assistance with massively parallel sequencing and access to the UPPMAX computational infrastructure.

This work is supported by Deutsche Forschungsgemeinschaft (DFG) through TUM International Graduate School of Science and Engineering (IGSSE), GSC 81.

The authors thank Thomas Walzthoeni for processing the sequencing data, and Malte Lücken, Elmar Spiegel, Ronan Le Gleut, and Giovanni Palla for discussions and valuable feedback. Furthermore, we thank Life Science Editors for revising the manuscript.

## Author contributions

### Conceptualization

AS, CH, KE, MPM, SE;

### methodology

AS, CH, AF, NB, JT, MM, MJ, ACP, ES; NGS;

### visualization

CH, SE, SF, AS;

### funding acquisition and supervision

KE, MPM, SE;

### project administration

AS, CH, AF;

### writing draft

KE, AS, CH, MPM, SE, AF;

### review and editing

MS, CSW, TB, FT

## Material and Methods

### Resource availability

#### Lead Contact

Further information and requests for resources and reagents should be directed to and will be fulfilled by the lead contact, Stefanie Eyerich

#### Data and Code Availability

Source code is available at github: https://github.com/Chillig/ST_biostatistical_analysis. RNA sequencing data can be obtained at GEO (www.ncbi.nlm.nih.gov/geo accession number: ….). This study did not generate new unique reagents.

#### Material Availability

This study did not generate new unique reagents

### Experimental Models and subject details

The study cohort consists of patients suffering from the non-communicable inflammatory skin diseases (ncISD) psoriasis (n=114), AD (n=52), lichen (n=35) and PRP (n=3). Gender was equally distributed in the cohort and mean age was 52,84 ± 17,48 years. Lesional and non-lesional skin samples were obtained for each disease. The study was approved by the local ethical committee (Klinikum Rechts der Isar, 44/16 S) and all patients gave written informed consent.

### Method Details

#### Spatial transcriptomics

##### Tissue sectioning, staining, library preparation

After obtaining non-lesional and lesional skin biopsies (6 mm), one third of each sample was immediately snap frozen in liquid nitrogen. Samples were then stored at −80°C until cryosectioning. Upon cryosectioning, samples were equilibrated to cryostat (NX70, Thermo Fisher Scientific) chamber temperature for at least 30 min and covered in optimal cutting temperature compound (OCT). Sections were taken at 10 μm thickness at −17°C and directly placed onto the Visium Spatial Gene Expression slide (10x Genomics). Slides were processed using the Visium Spatial Gene Expression Kit (10x Genomics) following the CG000239 Visium Spatial Gene Expression Reagent Kits - User Guide RevA. Optimal experiment conditions were investigated using the Visium Spatial Tissue Optimization Kit (10x Genomics) on independent healthy, lesional and non lesional skin samples, following the CG000238 Visium Spatial Gene Expression Reagent Kits - Tissue Optimization Rev A. To perform HE staining, samples were incubated in Mayer’s Hematoxylin (Dako) for 2 min and Eosin (Sigma) for 40 s, while Bluing buffer was omitted. Sections were permeabilized for 8 or 14 min and imaged using the Metafer Slide Scanning Platform (Metasystems) or the IX73 Inverted Microscope Platform (Olympus). Raw images were processed using VSlide software (Metasystems). Libraries of the individual data sets were pooled together separately and thereafter sequenced by the National Genomics Infrastructure (NGI, Sweden) on the Illumina NovaSeq platform using the recommended 28-10-10-120 cycle read setup.

### Sample annotation

HE images of corresponding samples were evaluated and annotated manually by two trained dermato-pathologists in a blinded manner using Loupe Browser (10x Genomics). Spots being present on tissue parts that were clearly destructed and broken off the section were marked and excluded from any further analysis. Samples were annotated for general morphology, anatomical structures, and specific cell types. Regarding general morphology, spots were categorized as “upper epidermis”, “middle epidermis”, “basal epidermis”, or “dermis”. Spots that were localised at the dermo epidermal junction were additionally marked as “interface”. To make the position of spots within the dermis comparable across the whole dataset, all spots categorized as “dermis” were further divided into “dermis 1” to “dermis 7” indicating the depth of the dermal layer in a standardized fashion.

### Data processing

52,020 spots were sequenced and samples were processed using 10x Visium Space Ranger-1.0.0. Quality control (QC) measures were applied on 64 samples with 56 passing QC. The sections were normalised and batch correction was applied to account for variances between the slides. DEG and pathway enrichment analysis were performed. Finally, the correlation between cytokine-secreting leukocytes and cytokine-dependent responder genes was investigated via a pseudo-bulk aggregation and a spatially weighted correlation approach. Due to acute inflammation, a high mitochondrial-fraction was anticipated, thus a conservative 25% cut-off was chosen. Spots with a minimum of 30 detected genes, and genes which were observed in at least 20 spots were considered. In addition, the QC enforced a minimum and maximum UMI-count of 50 and 500,000, respectively. The data were normalised using size factors calculated using the ‘scran’ R-package(*33*), log10 transformed, and a pseudo count of one was added to avoid log-transformation of zero(*34*). Highly variable genes were selected batch independently using ‘SCANPY’s’ highly_variable_gene function with flavor cellranger(*35*). The ST data set was batch corrected with ‘scanorama’(*36*) accounting for the variances between the slides. Further, the data set was dimensionally reduced by applying a principal component (PC) analysis with n_pcs=8 and embedded in a neighborhood graph with n_neighbors=15. Subsequently, the data were represented in a 2D UMAP plot.

#### Clustering of transcriptomes

The ST analysis benefited from expert annotations of dermato-pathologists, thus forming the clusters based on epidermis layers, interface and dermis depths 1-7. For the clustering of the scRNA-seq data, we leveraged the Leiden algorithm and determined the number of clusters by the maximum silhouette score, and prior knowledge, i.e. enriched marker genes in stable clusters. At a resolution of 0.1, the maximum silhouette score was 0.54.

#### Spatial enrichment of cytokines in specific skin layers

Unnormalised count matrices, and a targeted analysis scrutinised for *IL17A*, *IFNG*, and *IL13* was used to analyse cytokine expression compared to a housekeeping gene, *GAPDH*, in ST. Cytokine expression levels were quantified within the manually curated skin layers, and significant spatial enrichments were tested with Wilcoxon signed-rank test.

#### Differential gene expression (DEG) and pathway enrichment analysis

To characterize cytokine expressing cells, leukocytes were defined by the marker genes *CD2*, *CD3D*, *CD3E*, *CD3G*, *CD247*, and *PTPRC* in the ST and single-cell datasets. Leukocytes were defined as cytokine-positive if at least one UMI-count of the cytokine gene was detected. Prior to the DGE analysis, the counts were normalised using size factors calculated on the whole data set.

Genes characterising cytokine-positive spots were compared with cytokine-negative spots to obtain differentially expressed genes on a spot-level using ‘glmGamPoi’(*37*) and the multiple testing method ‘Benjamini-Hochberg’ (BH). In addition to the unnormalised counts, the calculated size factors were provided and biological variances were included as fixed effects in the design matrix. In the design matrix the covariates cellular detection rate (cdr), patient, and annotation were included. This enabled to account for variances between the fraction of genes being transcribed in a cell(*38*), and the difference in gene expression between cells that are located in different tissue types and are of different cell types, respectively. The following model was used for the ST data set

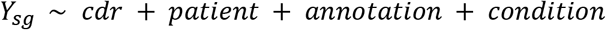

or for the single psoriasis patient scRNA-seq data set

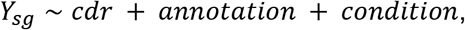

where *Υ_sg_* is the raw count of gene *g* in the cell or spot *s*. A gene is called significantly differentially expressed if it meets the cut-off parameters of p-value <= 0.05 and |log2FC| >=1. Pathway enrichment analysis was performed using the Bioconductor packages ‘ReactomePA’(*39*) and ‘org.Hs.eg.db’(*40*) and illustrated using the Bioconductor package ‘enrichplot’(*41*). The p-values of the pathways were corrected using the BH method and a p-value and q-value cut-off of 0.05 was applied.

#### Correlation between cytokines and responder genes

ST spots were annotated either as cytokine-positive, responder-positive, or other. Spots that contained both cytokine and responder genes were labelled as cytokine-positive. Spots containing neither a cytokine nor a responder gene were labelled as other. As the responder gene signature was obtained from *in vitro* stimulated primary human keratinocytes experiment, the correlation analysis focused solely on the epidermis.

To describe the spatial relationship between cytokine-positive and responder-positive spots, we developed a density-based clustering method that leverages confirmed cytokine-positive spots as seeds. Our method provides the possibility to cluster cytokine-positive spots based on three conditions. First, cytokine-positive transcript points were connected if they were in the neighbourhood of a unit circle resulting in a cytokine graph. Second, a cluster was built by adding responder spots to the graph if they were among the nearest neighbour spots in the vicinity of the unit circle. Third, clusters were merged if they shared responder positive capturing points. By applying these conditions, the clusters were characterized based on the density of cytokine-positive spots and the response close to them. Accordingly, based on these conditional density clusters a weighted Pearson correlation was calculated. The weights were determined by the number of cytokine-positive spots.

In more detail, the adjacent cytokine-positive locations were obtained per sample using the KDTree algorithm(*42*) with the Euclidean metric and a maximum distance of 2.0. For this purpose, the index array provided by 10X Genomics was used. Afterwards, the locations of adjacent cytokine-positive mRNA capturing points were connected using a graph as backbone. Here, the nodes were the cytokine-positive spots and the edges equal the distance between the spots. Moreover, the nearest neighbor responder spots were determined by

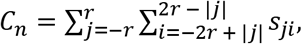

where *r* is the radius of the cluster and *s_ji_* is the nearest neighbor spot in row *j* and column *i*. Then the cytokine-positive graph was merged together with the nearest neighbor responder spots resulting in an agglomerated graph. Finally, the counts of responder genes and cytokines in each cluster were read out and a weighted Pearson correlation was calculated. The weights were determined by the number of cytokine-positive spots in the graph to account for the size and impact of the density cluster.

#### In situ hybridization

*In situ* hybridization was performed using the RNAScope® Multiplex Fluorescent V2 Assay for paraffin embedded tissue sections (Advanced Cell Diagnostics, Newark, CA) on lesional skin sections of psoriasis, AD, and lichen (5 μm each). The assay was performed using probes designed by ACD targeting human *IL17A, IFNG* or *IL13* mRNA. Positive control sections were prepared using human peptidylprolyl isomerase B (PPIB) probe whereas negative controls were assessed using bacterial gene probes. Briefly, target probes were hybridized followed by signal amplification according to manufacturer’s protocol. Each probe was stained by Opal 690 (Akoya Biosciences, Marlborough, MA) using a single-plex setup. Subsequently, skin sections were examined using microscope slide scanner (Axio Scan.Z1 Zeiss, Germany) at 20x magnification. Then, images were visualized using QuPath software(*43*). Images were individually evaluated by two trained dermato-pathologists in a blinded manner. Cells were counted positive if punctate-dot RNAscope® signal co-localized with nuclear staining.

## Supplemental Information

### Material and Methods

#### Immunohistochemistry

5 μm sections of paraffin embedded skin samples were air-dried overnight at 37 °C, dewaxed and rehydrated. Stainings were performed by an automated BOND system (Leica) according to the manufacturer’s instructions: epitope retrieval was performed at pH6 in epitope retrieval solution (DAKO) and incubated with goat anti-human IL-17A (R&D Systems) followed by a biotinylated anti-goat secondary antibody (Vector Laboratories). For detection of specific binding, streptavidin peroxidase and its substrate 3-amino-9-ethyl-carbazole (DAKO) were used. All slides were counter stained with hematoxylin. Stainings without primary antibodies were used as negative control. Positive cells were counted in four to nine visual fields per condition.

#### Isolation of primary human T cells and *in vitro* stimulation

Peripheral blood mononuclear cells were isolated from peripheral blood of healthy donors by density centrifugation. Primary human Pan T cells were then isolated using magnetic beads (Pan T cell isolation kit, Miltenyi Biotec), followed by CD4 (human CD4 microbeads, Miltenyi Biotech) or CD8 (human CD8 microbeads, Miltenyi Biotec) isolation. Defined numbers of cells were stimulated with platebound anti-CD3 and anti-CD28 antibodies (0.75 μg/ml; BD Biosciences) for 10 min, 1 h or 6 h, or were left unstimulated. Stimulated T cells were collected after 10 min, 30 min, 1 h, 6 h, 12 h, or 24 h stimulation and RNA was isolated for subsequent real time PCR analysis with the following primers: IL-17A (fw: CAATCCCCAGTTGATTGGAA; rev: CTCAGCAGCAGTAGCAGTGACA, IFN-γ (fw: TCAGCCATCACTTGGATGAG; rev: CGAGATGACTTCGAAAAGCTG), IL-13 (fw: TGACAGCTGGCATGTACTGTG; rev: GGGTCTTCTCGATGGCACTG), 18S (fw: GTAACCCGTTGAACCCCATT; rev: CCATCCAATCGGTAGTAGCG).

#### Flow cytometry of skin T cells

Primary human T cells (n=52) were isolated by digestion of fresh human skin biopsies (Ø 6 mm) in RPMI containing FCS, Collagenase type IV (Worthington), and Deoxyribonuclease I (Sigma) at 37°C overnight followed by dissociation using the gentleMACS Dissociator (Miltenyi Biotec). Freshly isolated skin T cells were passed over a cell strainer and directly used for flow cytometric analysis. For flow cytometric analysis, T cells were stimulated with PMA/Ionomycin (10 ng/ml and 1 μg/ml, respectively) (both Sigma) for 5 h in the presence of Brefeldin A and Monensin (both BD Biosciences). Surface staining was performed at 4°C and followed by fixation/ permeabilization using the fixation/permeabilization kit (BD Biosciences). Staining of intracellular cytokines was performed at room temperature. Antibodies used were CD3-Bv650, CD4-BV421, CD8-APCCy7 (BD Biosciences), IL-17A-PeCy7, IFN-γ-PerCPCy5.5, TNF-α-BV510 (BioLegend), IL-22-Pe (eBioscience), IL-10-APC (Miltenyi Biotec).

#### Single-cell RNA sequencing

A lesional skin sample (6 mm) was taken from a psoriasis patient and digested immediately for 3 h at 37 °C using the MACS whole skin tissue dissociation kit (Miltenyi Biotec) and the gentleMACS Dissociator (Miltenyi Biotec) according to manufacturer’s protocol. The obtained cells were stained for CD3 (Biolegend, 300450) and CD45 (BD Biosciences, 563880) and sorted using a FACSAria Fusion (BD). Here, dead cells and doublets were gated out and cells sorted based on size (FSC/SSC) and CD3/ CD45 expression into three populations: skin cells (keratinocytes), T cells (CD45+, CD3+), and APCs (CD45+, CD3−). The obtained cells were mixed in equal ratio (1:1:1) to a final cell number of 16,000 and used as input for the sc library generation by the 10x Genomics kit (Chromium Single Cell 3’ Kit v3) according to the manufacturer’s protocol. The libraries were sequenced on an Illumina HiSeq4000 via paired-ends with a read length of 2 x 150 bp at a sequencing depth of 40 million reads.

##### scRNA-seq data processing

The pre-processing and QC of the scRNA-seq data was identical to ST, besides enforcing a minimum of 500 genes per cell, and a minimum and maximum UMI-count of 600 and 25,000, respectively. In addition, according to the scrublet pipeline(*44*), no doublets were detected. Additionally, no batch effect was detected in the scRNA-seq data and the number of PCs was set to n_pcs = 7.

## Supplemental Figures

**Figure S1:**
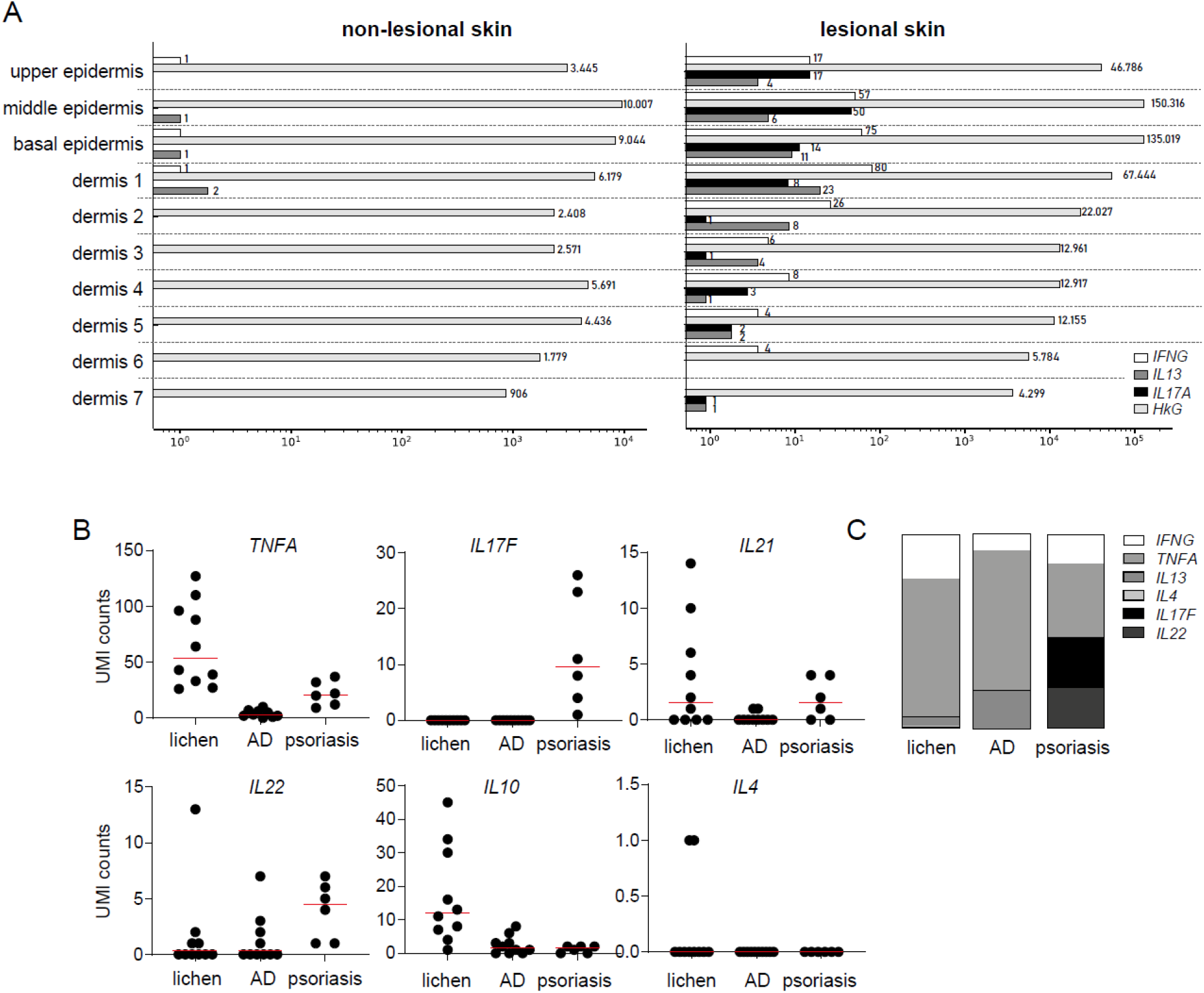
NcISD are characterized by low cytokine UMI counts in skin. **A)** Total UMI counts in spatial sections for *IFNG, IL17A, IL13* and the housekeeping gene *GAPDH* (*HkG*) in non-lesional and lesional skin separated by the location in the skin. **B)** UMI counts for selected cytokines in sections of lichen (n=10), atopic dermatitis (AD (n=10), and psoriasis (n=6). **C)** Percentage of disease relevant cytokine UMI counts in lichen, AD, and psoriasis normalised to 100%.

**Figure S2:**
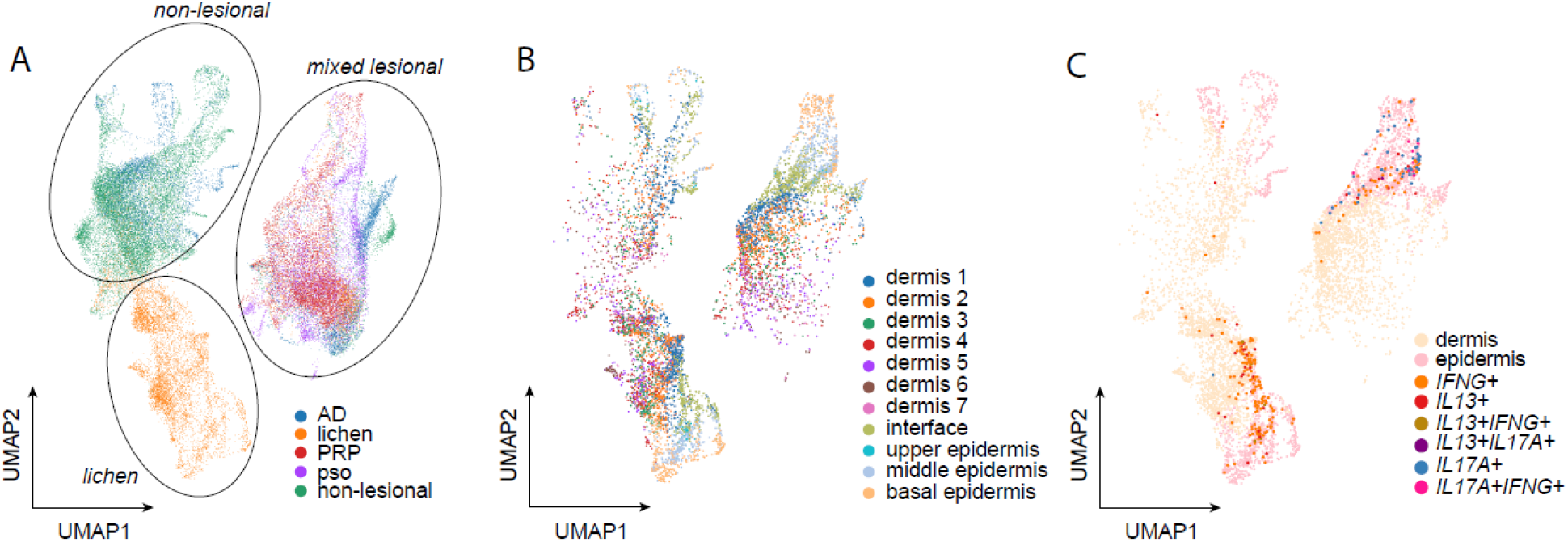
Cytokine positive spots are located in the epidermis and enriched in lesional skin. **A)** UMAP plot highlighting the origin of each spatial spot according to the lesional and non-lesional skin as well as disease (psoriasis (Pso), atopic dermatitis (AD), lichen planus (lichen), and pytiriasis rubra pilaris (PRP)). **B)** UMAP highlighting the manually annotated tissue layers basal, middle and upper epidermis and dermis 1-7 in all spatial samples expressing leukocyte markers (n=56). **C)** UMAP plot indicating cytokine producing cells in epidermis and dermis in spatial sections expressing leukocyte markers (n=56).

**Figure S3:**
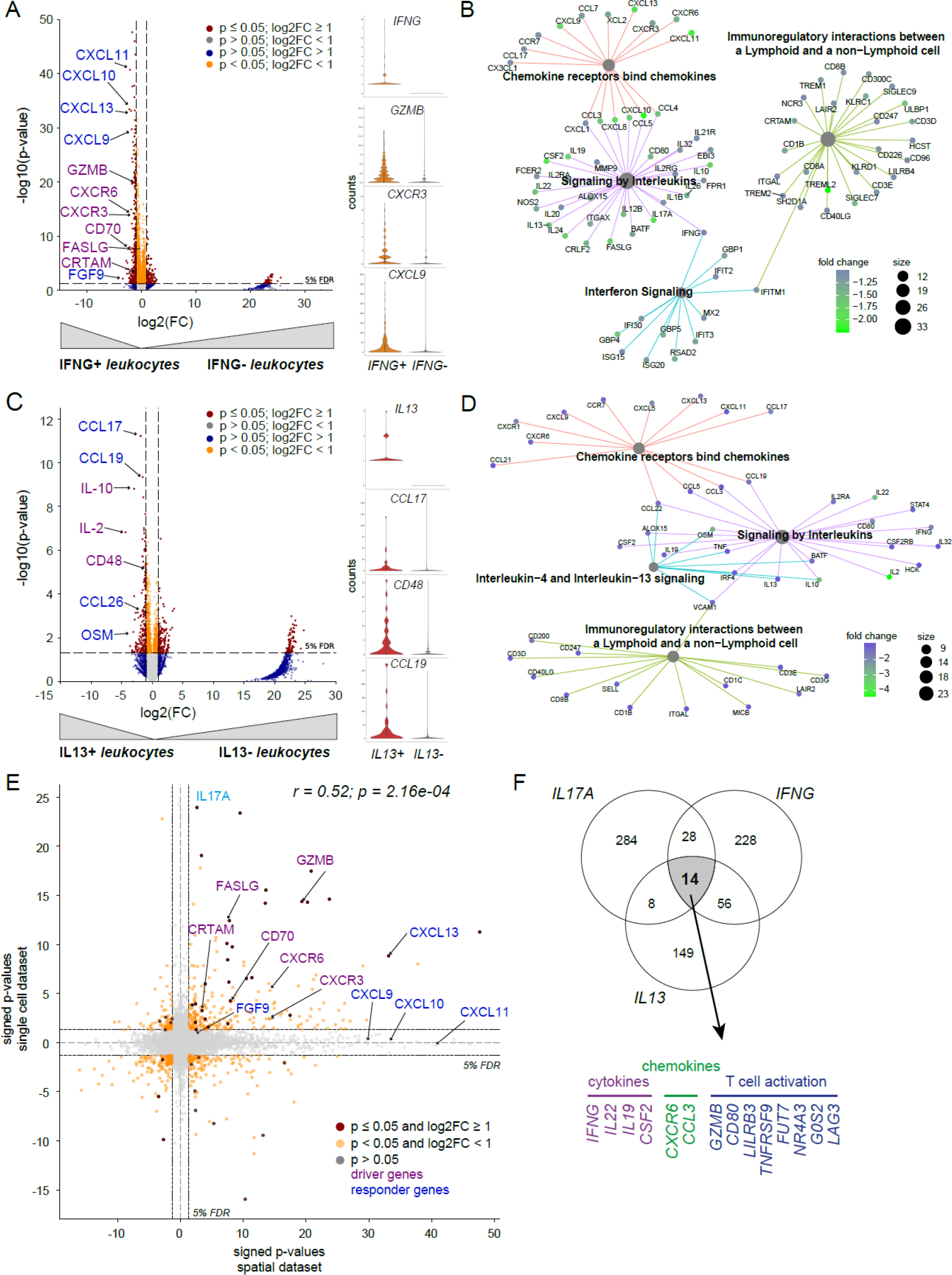
Cytokine positive spots are characterized by specific gene signatures. **A)** Volcano plot analysing the gene expression profile of *IFNG* positive (*IFNG*+) *versus IFNG* negative (*IFNG*−) cells in ST sections (n=56). Coordinates for *IFNG* (−37.4/255) are not shown. Violin plots show expression of selected genes in both groups. **B)** Gene set enrichment analysis of genes co-expressed with *IFNG*. **C)** Volcano plot analysing the gene expression profile of *IL13* positive (*IL13*+) *versus IL13* negative (*IL13*−) cells. Coordinates for *IL13* (−38.1/69.7) are not shown. Violin plots show expression of selected genes in both groups. **D)** Gene set enrichment analysis of genes co-expressed with *IL13*. **E)** Comparison of the DEG analysis of *IFNG* positive spatial spots and *IFNG* positive single cells in a signed p-value plot. Significantly expressed genes in both data sets are shown in red. Purple and blue labeling indicates T cell derived genes and skin response genes, respectively. **F)** Common genes shared between all cytokine positive spatial spots.

**Figure S4:**
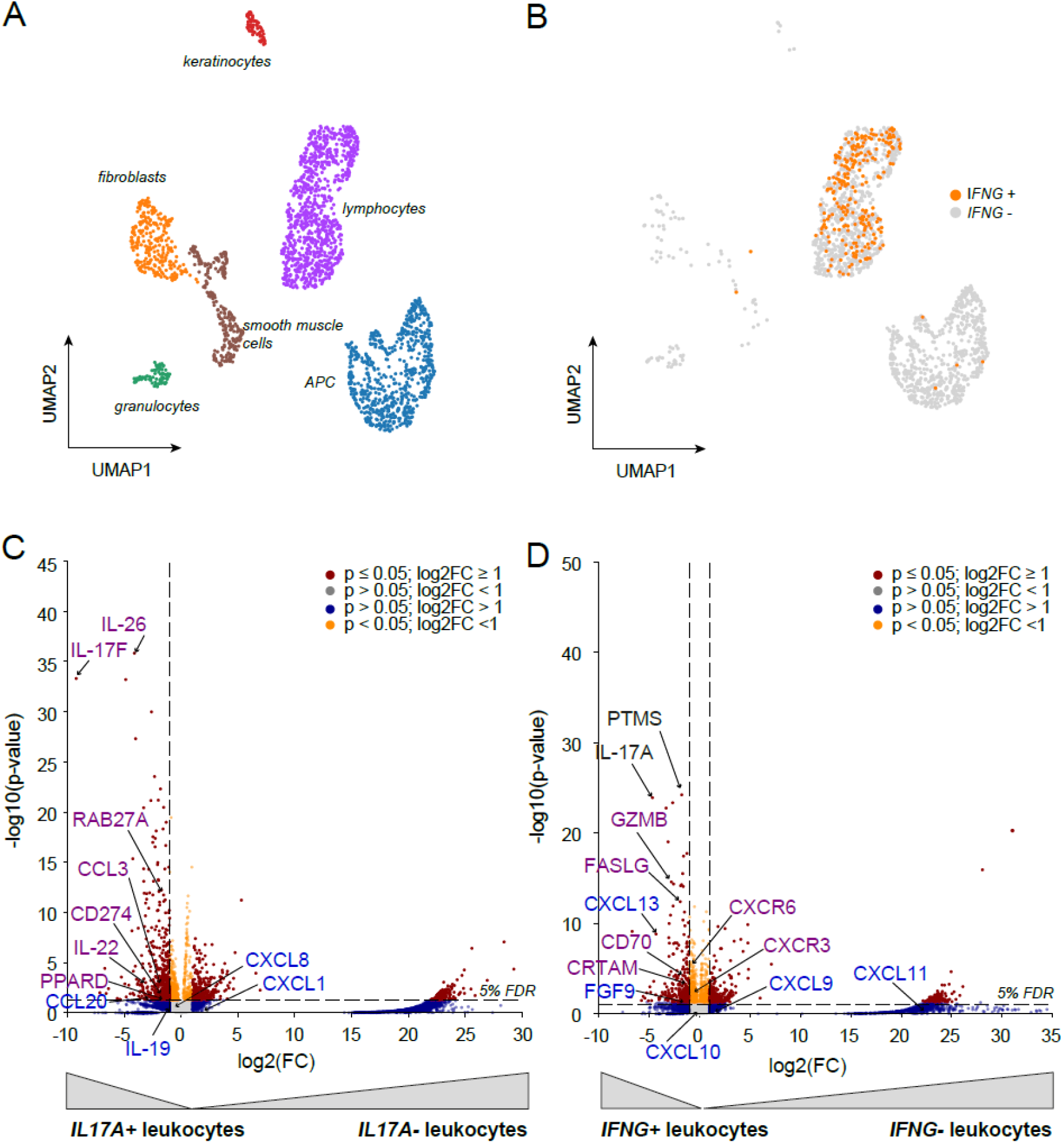
Single cell analysis reveals specific gene signatures for IL-17A and IFN-γ expressing cells. **A)** UMAP plot indicating the composition of cellular clusters in the single cell RNASeq dataset. **B)** UMAP plot highlighting *IFNG* positive (*IFNG*+) leukocytes in orange and *IFNG* negative (*IFNG*−) leukocytes in grey. **C)** Volcano plot analysing differentially expressed genes (DEG) in *IL17A* positive (*IL17A*+) versus *IL17A* negative (*IL17A*−) leukocytes or **D)** *IFNG* positive (*IFNG+) versus IFNG* negative (*IFNG*−) leukocytes in the single cell data set. Coordinates of *IL17A* (−36.9/256) and *IFNG (−*36/263) are not shown.

**Figure S5:**
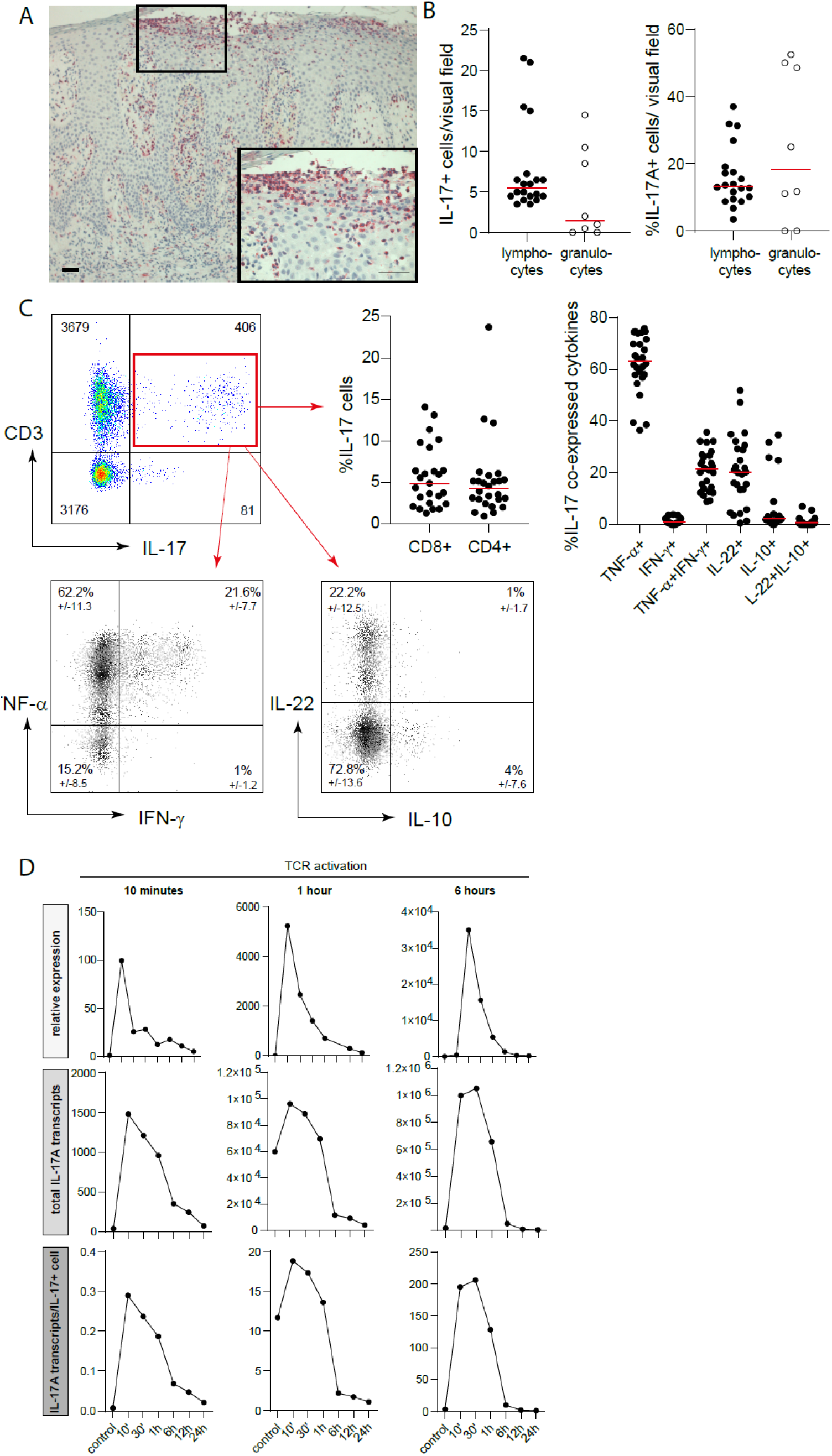
IL-17A expression in skin lesions and infiltrating T cells and its short half-life time and low copy numbers in in vitro stimulated T cells. **A)** Representative staining of IL-17A by immunohistochemistry in a psoriasis section. **B)** Number (left panel) and percentage (right panel) of IL-17A+ lymphocytes and granulocytes per visual field in psoriasis sections stained by immunohistochemistry (n=20 patients). **C)** Representative flow cytometry staining of T cells derived from lesional psoriasis skin. CD3+IL-17A+ T cells were analysed for co-production of IL-22, TNF-α, IL-10, and IFN-γ by intracellular flow cytometry. The graphs indicate the percentage of CD4+ and CD8+ cells amongst the CD3+IL-17A+ cells and the frequency of IL-17A producing cells co-expressing one or two other cytokines (n=52). **D)** CD4+ T cells were isolated from blood of healthy donors and stimulated with anti-CD3/anti-CD28 antibodies (TCR activation) for the indicated time. RNA was isolated over a time course of 24 h and analysed for the expression of IL-17A by real time PCR. Relative expression of IL-17A was calculated to unstimulated cells (upper panel). Total transcript numbers of IL-17A were determined in each stimulatory condition using a standard curve (middle panel). By dividing the total transcript numbers by the number of cells per stimulatory conditions, the transcript number per cell could be identified (lower panel).

**Figure S6:**
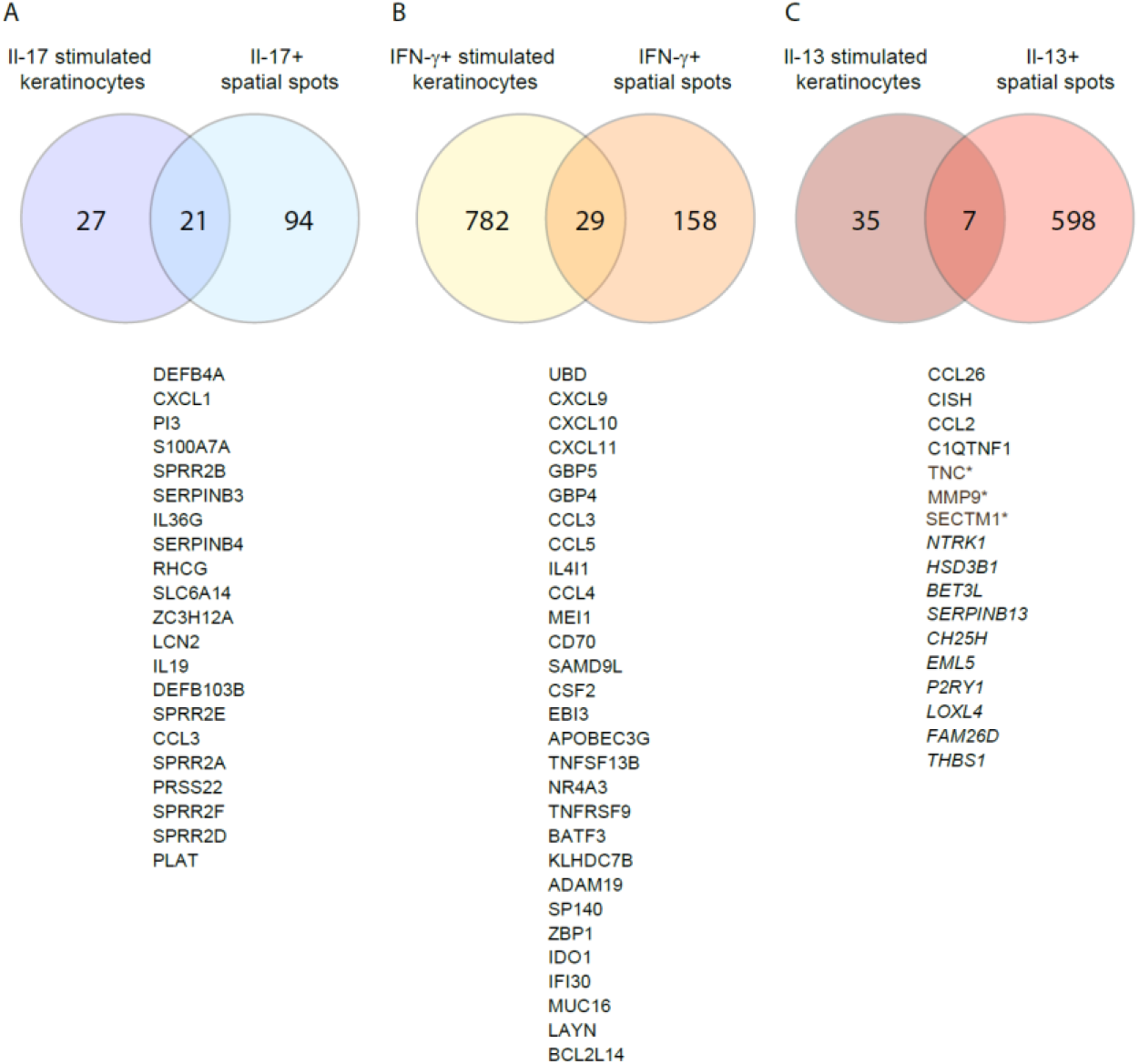
Identification of cytokine responder genes for spatial correlation. Primary human keratinocytes were stimulated in 2D cultures with recombinant IL-17A, IFN-γ or IL-4/IL-13 (20 ng/ml each) for 16 h. Total RNA was isolated and whole genome expression arrays (SurePrint G3 Human GE 8X60K BeadChip (Agilent Technologies)) were performed according to the manufacturer’s instructions. Gene expression data was filtered for p-value <0.05, adjusted p-value <0.05, and log2 FC >1.5 for **A)** *IL17A* and **B)** *IFNG* or log2FC >1 for **C)** IL-13. Differential genes co-expressed in spatial spots with **A)** *IL17A*, **B)** *IFNG*, or **C)** IL-13 were filtered for p-value <0.05, adjusted p-value <0.05, and log2 FC >1.5. Gene expression lists were compared by Venn diagram analysis and commonly expressed genes were selected as cytokine specific responder genes for the indicated cytokine. For *IL13* only 7 genes were commonly expressed with 3 genes (*SECTM1, TNC, MMP9*) (*) being also expressed in IFN-γ stimulated keratinocytes. These genes were removed and genes in italic added according to literature analysis.

**Figure S7:**
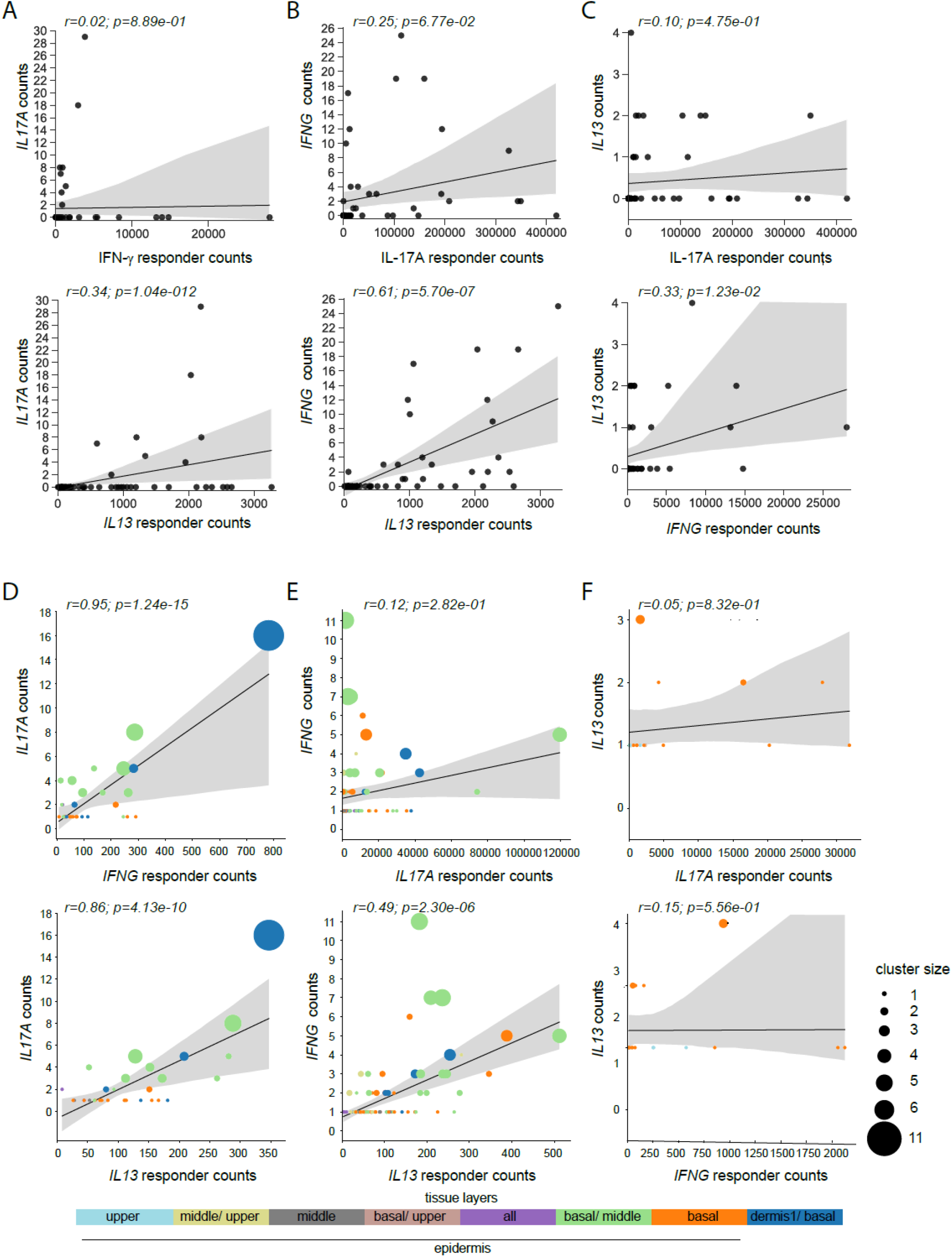
Cytokine UMI counts only correlate with their specific responder genes, but not those of other cytokines. **A-C)** Pearson correlations between the sum of counts of cytokines and permuted cytokine responder genes. Each point represents a tissue sample. Correlation between **A)** *IL17A* and responder genes of *IFNG* and *IL13*, **B)** *IFNG* and responder genes of *IL17A* and *IL13*, and **C)** *IL13* against *IL17A* and *IFNG* response signatures. **D-F)** Weighted spatial correlation incorporating the spatial relation of cytokines and the permuted response located in the epidermis. Each point in the plots represents the sum of the counts of cytokines and responders in a cluster and the size of each point. **D)** correlation of *IL17A* and responder genes of *IFNG* and *IL13*, **E)** *IFNG* and responder genes of *IL17A* and *IL13*, and **F)** *IL13* against *IL17A* and *IFNG* response signatures.

